# Phylogenetic conservation of bacterial environmental responses predicts soil bacterial biogeographic patterns

**DOI:** 10.64898/2026.08.03.742428

**Authors:** Mingming Xia, Kazuo Isobe, Jennifer BH Martiny

**Affiliations:** Institute of Ecology, State Key Laboratory of Vegetation Structure, Function and Construction, and College of Urban and Environmental Science, Peking University, Beijing, China; Department of Ecology and Evolutionary Biology, University of California, Irvine, CA, USA

**Keywords:** soil bacterial biogeography, phylogenetic trait conservation, community turnover, pH gradient, community prediction

## Abstract

Soil bacterial communities exhibit biogeographic patterns along environmental gradients, yet why some environmental factors contribute more strongly to community turnover than others remains poorly understood. Here, we tested whether this variation can be explained by the phylogenetic depth at which bacterial responses to each environmental factor are conserved. Across 40 forest sites in Japan spanning multiple soil and climatic gradients, environmental factors whose bacterial responses were conserved at deeper phylogenetic levels contributed more strongly to bacterial community turnover. We further asked whether phylogenetic clades that share similar environmental responses represent ecologically meaningful units for understanding bacterial community responses. Using soil pH as a focal test case, we found that response-defined clades improved prediction of taxon-level abundance shifts and community-level compositional shifts compared with models that treated taxa as independent units. Together, these findings show that the phylogenetic depth of bacterial environmental responses links trait conservation, community turnover and soil bacterial biogeographic patterns.

**Significance Statement:** Soil bacterial communities form biogeographic patterns along environmental gradients, but it remains unclear why some environmental factors drive stronger community turnover than others. This study shows that the strength of bacterial community turnover can be predicted from the phylogenetic depth at which bacterial responses to environmental factors are conserved. Across forest soils in Japan, deeply conserved bacterial responses were linked to stronger community turnover, and clades sharing conserved responses improved prediction of both taxon- and community-level shifts. These findings identify phylogenetically conserved response structure as an organizing principle for understanding and predicting soil bacterial biogeographic patterns.

## Introduction

Soil bacterial community composition varies along environmental gradients and exhibits clear biogeographic patterns^1^. Although associations between environmental factors and bacterial community composition have been extensively documented^2^, why some environmental factors exert stronger influence on community turnover than others remains poorly understood. Resolving this question is essential for moving beyond the description of bacterial biogeographic patterns toward a mechanistic understanding of how environmental factors organize bacterial communities.

Bacterial responses to environmental conditions are often phylogenetically conserved, such that closely related taxa tend to respond in similar ways to environmental change^3–8^. Such conserved responses are thought to arise because related bacterial taxa share functional traits and physiological systems that shape how they respond to environmental conditions. Importantly, however, the phylogenetic depth at which bacterial responses are conserved differs among environmental factors^7,8^; responses to some factors are shared across deeper phylogenetic lineages, whereas responses to other factors are conserved only among more closely related taxa. This variation in conservation depth is expected to reflect the physiological complexity and integration of the traits underlying each response. Responses governed by complex, multi-component systems should be conserved at deeper phylogenetic levels, whereas those based on simpler or more modular functions should be conserved more shallowly^9^.

This variation in the phylogenetic depth at which bacterial responses are conserved leads to two linked hypotheses for bacterial biogeography. First, environmental factors whose bacterial responses are conserved at deeper phylogenetic levels should drive compositional turnover across broader phylogenetic scales, involving shifts among distantly related clades^10^ (Fig. 1A). In contrast, factors whose responses are conserved only shallowly should generate more limited compositional shifts, primarily within closely related clades. This pattern has been demonstrated within the marine cyanobacterial lineage *Prochlorococcus*, where light intensity, whose response is deeply conserved, structures major clades; water temperature, whose response is conserved at intermediate phylogenetic depth, explains variation at intermediate phylogenetic scales; and nutrient availability, whose response is more shallowly conserved, drives fine-scale diversification^11,12^.

**Fig. 1.**
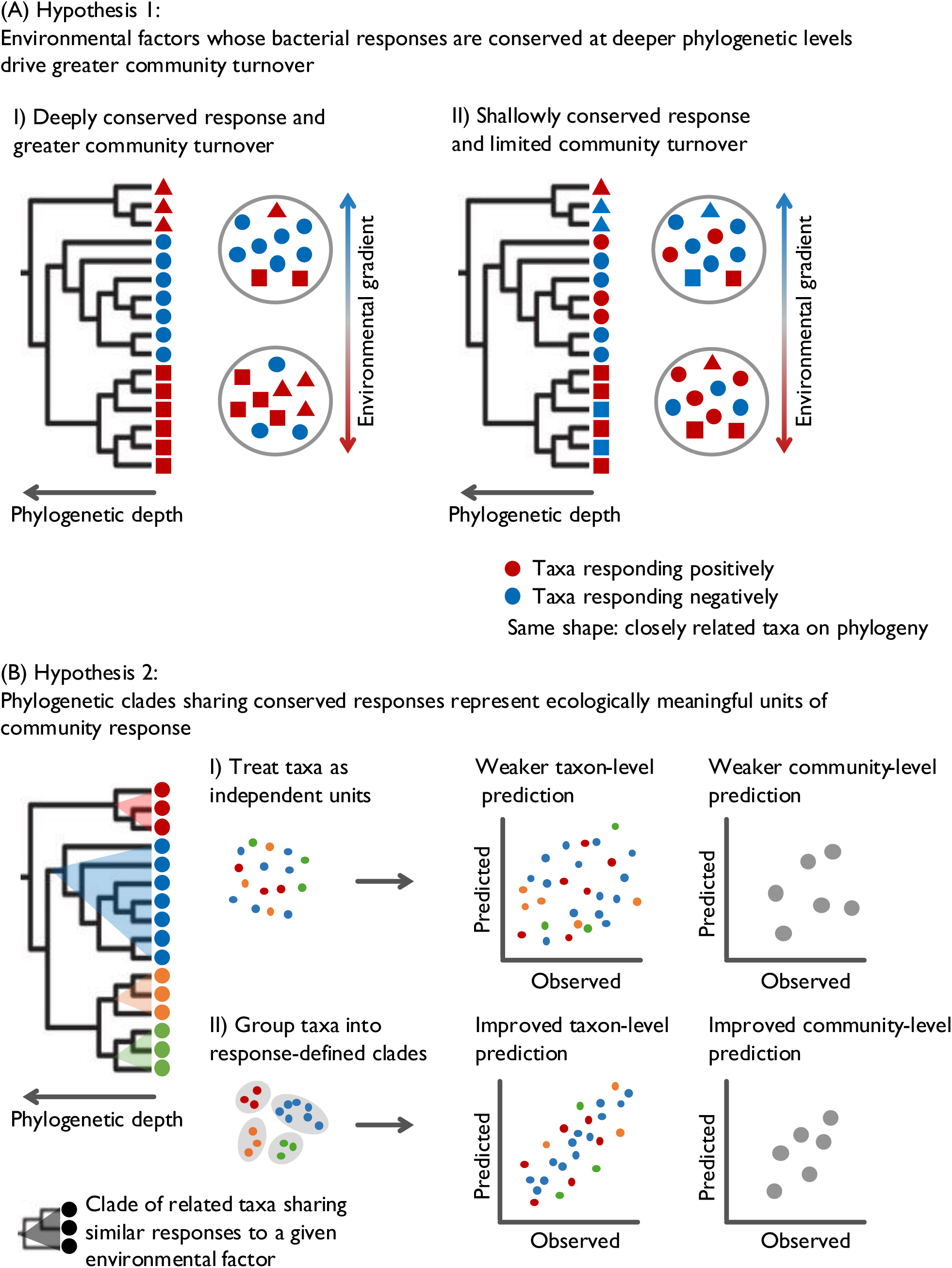
Conceptual framework linking phylogenetic response conservation to bacterial biogeography. (A) Environmental factors whose bacterial responses are conserved at deeper phylogenetic levels are expected to drive stronger community turnover by organizing compositional shifts across broader phylogenetic scales. In contrast, factors whose responses are conserved only shallowly are expected to generate more limited turnover, primarily among closely related taxa. Symbol colour indicates response direction, and symbol shape indicates phylogenetic position; taxa with the same shape are closely related. (B) If phylogenetically conserved responses organize bacterial community shifts, then the clades in which those responses are conserved should represent ecologically meaningful units of community organization. Prediction provides an operational test of this idea: response-defined clades should improve prediction of taxon-level abundance shifts compared with models that treat taxa as independent units. Stronger improvements are expected where shared response tendencies persist across greater phylogenetic distances, as illustrated by the blue-highlighted clade containing taxa with similar responses, and improved taxon-level prediction should translate into improved prediction of community-level compositional shifts.

Second, if this primary hypothesis is correct, then the clades in which bacterial responses are conserved should represent ecologically meaningful units of community organization rather than arbitrary phylogenetic groupings (Fig. 1B). Such response-conserved clades should capture shared response information that is lost when taxa are treated as independent units. Prediction provides an operational test of this idea: ecologically meaningful response-conserved clades should improve prediction of taxon-level abundance shifts, with stronger improvement expected where shared response tendencies persist across greater phylogenetic distances. If these clades capture ecologically relevant response structure, improved taxon-level prediction should also scale up to better prediction of community-level compositional shifts.

Here, we extended this trait-based phylogenetic framework to whole soil bacterial communities by analyzing bacterial communities from 40 forest sites across Japan (Fig. S1). We first tested whether environmental factors whose bacterial responses are conserved at deeper phylogenetic levels contribute more strongly to compositional turnover across whole soil bacterial communities (Fig. 1A). We then asked whether the clades in which such responses are conserved function as ecologically meaningful units of community organization (Fig. 1B). Because soil pH showed both the strongest association with community turnover and the deepest phylogenetic conservation of bacterial responses, we used pH as a focal test case. Using prediction as an operational test, we examined whether pH response-defined clades improved prediction of taxon-level abundance shifts, whether this improvement was greater where shared response tendencies persisted across greater phylogenetic distances, and whether improved taxon-level prediction translated into improved prediction of community compositional shifts across ecosystems. Together, our results suggest that the phylogenetic depth of bacterial environmental responses provides a mechanistic link between trait conservation, community turnover and bacterial biogeographic patterns across ecosystems.

## Results

### Phylogenetic depth of conserved responses explains variation in bacterial community turnover

To test our first hypothesis, we asked whether environmental factors whose bacterial responses are conserved at deeper phylogenetic levels contribute more strongly to compositional turnover across soil bacterial communities. We first quantified the relative contributions of soil pH, soil total carbon (TC), carbon-to-nitrogen ratio (C/N), mean annual temperature (MAT), and mean annual precipitation (MAP) to variation in bacterial community composition using generalized dissimilarity modelling (GDM). Together, these variables explained 57.6% of the total compositional variation in bacterial communities, with soil pH contributing the most (26.2%), followed by C/N (14.4%), MAT (11.9%), TC (3.2%) and MAP (2.0%) (Fig. 2A).

**Fig. 2.**
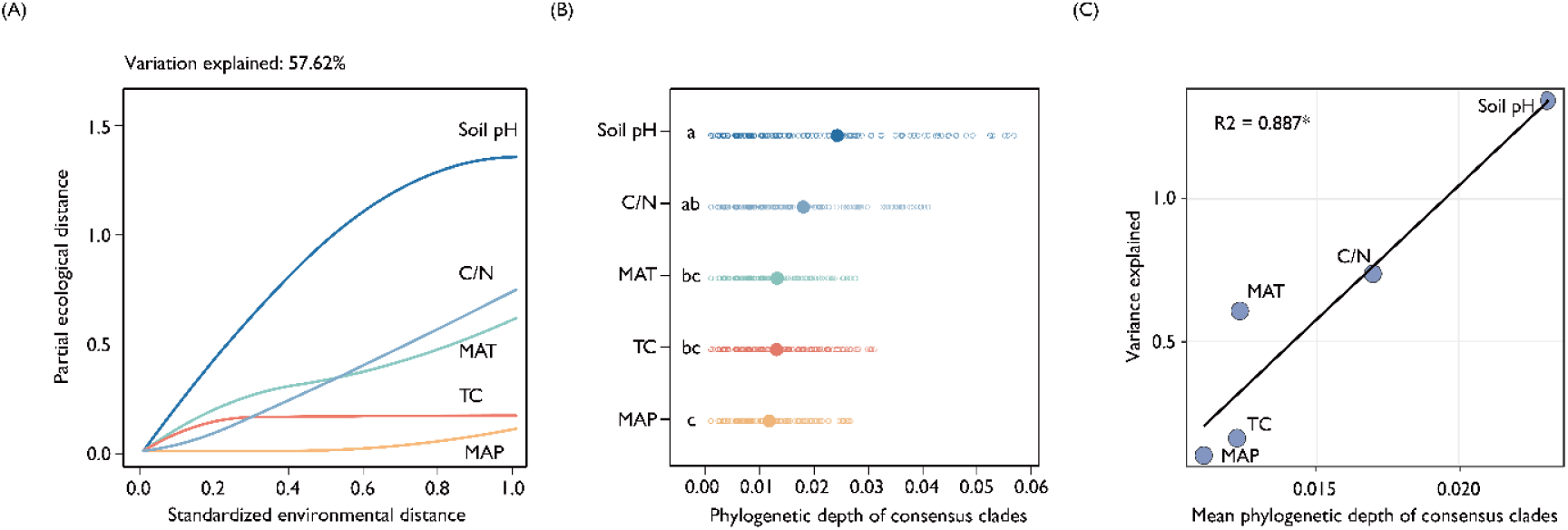
Phylogenetic conservation of bacterial responses to environmental factors and its relationship to community turnover. Environmental predictors include soil pH, soil total carbon (TC), C:N ratio (C/N), mean annual temperature (MAT), and mean annual precipitation (MAP). (A) Generalized dissimilarity model (GDM) of microbial community turnover along environmental gradients. Fitted I-spline curves show how microbial β-diversity changes with standardized differences in each environmental predictor while holding others constant. The x-axis denotes standardized environmental distance (0–1), and the y-axis denotes the partial ecological distance attributable to each variable. Curve height reflects the total amount of community turnover explained by each variable, whereas curve slope indicates the rate of turnover along the gradient. The full GDM explains 57.6% of the variation in community composition. (B) Distributions of the phylogenetic depth of response-consensus clades, defined as clades in which ≥90% of descendant ASVs exhibit a significant response (Spearman correlation, p < 0.05). Phylogenetic depth was estimated using the consenTRAIT algorithm. Open circles represent individual consensus clades, and filled circles represent the mean depth across clades (τD). Values of τD are provided in Table S1. Different letters indicate significant differences in depth among environmental variables (one-way ANOVA followed by Tukey’s HSD test, p < 0.05). (C) Relationship between the mean phylogenetic depth (τD) of response-consensus clades and the contribution of each environmental factor to bacterial community turnover.

We next quantified the phylogenetic depth of conserved responses to each environmental factor using the consenTRAIT framework. In this analysis, response-consensus clades were defined as phylogenetic clades in which ≥90% of descendant ASVs exhibited significant responses to a given environmental variable, and the mean phylogenetic depth of these clades (τD) was calculated for each variable. We use the term response-consensus clades specifically for the consenTRAIT-defined clades used to estimate τD, to distinguish them from the response-defined clades used below in the pH prediction analyses. The degree of phylogenetic conservation varied substantially among environmental factors. Soil pH exhibited the greatest mean phylogenetic depth across response-consensus clades (τD = 0.023, corresponding to approximately 4.6% sequence dissimilarity among ASVs), followed by C/N (τD = 0.017), whereas MAT, TC and MAP showed lower τD values (0.011–0.013) (Fig. 2B; Table S1). Notably, soil pH and C/N showed significant phylogenetic conservation across all response categories, including all, positive and negative responses (Fig. S2; Table S1). Consistent with these patterns, pairwise comparisons among ASVs showed that differences in response strength increased with phylogenetic distance for soil pH and C/N, whereas this pattern was not evident for MAT, TC or MAP (Mantel tests; Fig. S3). Given the particularly deep and consistent conservation of soil pH responses, we further confirmed that pH responses were phylogenetically conserved within major bacterial phyla, although differences in τD among phyla should be interpreted cautiously because they may reflect ASV number and phylogenetic sampling density (Table S2).

Finally, we tested whether the phylogenetic depth of conserved responses predicted the contribution of each environmental factor to bacterial community turnover. We found a strong positive relationship between τD and the GDM-estimated contribution of each environmental factor (R² = 0.887, p = 0.017; Fig. 2C), indicating that environmental factors with more deeply conserved bacterial responses contributed more strongly to bacterial community turnover. When responses were separated by direction, this relationship was significant for positive responses (R² = 0.894, p = 0.015; Fig. S4A) but weak for negative responses (R² = 0.342, p = 0.299; Fig. S4B). These results support our first hypothesis that the phylogenetic depth of conserved responses helps explain why some environmental factors are associated with stronger bacterial community turnover than others.

### Response-defined clades capture shared structure in pH-driven ASV abundance shifts

To test our second hypothesis, we focused on soil pH as the strongest empirical test case for our framework, because it showed both the deepest phylogenetic conservation of responses and the largest contribution to community turnover. We first identified 59 response-defined clades associated with soil pH, grouping related ASVs that shared broad pH-response tendencies (PhyloFactor analysis; Fig. 3A). However, an overall positive or negative response can mask nonlinear pH-response patterns and finer-scale variation among ASVs within the same clade. We therefore quantified range-specific ASV-level responses within these clades across the pH gradient, including the pH intervals over which each ASV showed a consistent abundance response and the magnitude of that response (TITAN2 analysis; Fig. S5). This combined approach allowed the prediction model to incorporate both clade-level response structure and range-specific ASV-level variation. By contrast, pH optima, defined as the pH at which relative abundance was maximized within the sampled range, varied substantially among ASVs within the same response-defined clade (Fig. 3B). We therefore did not include pH optimum as a predictor in the model.

**Fig. 3.**
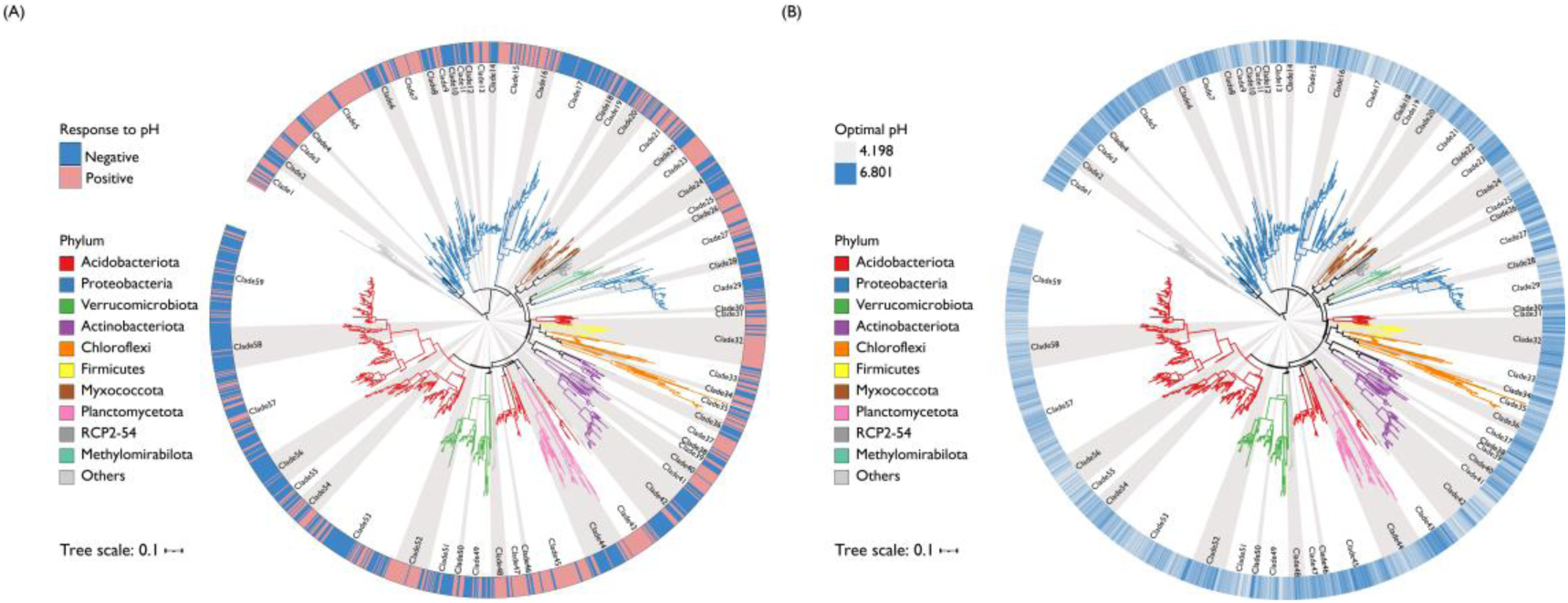
Phylogenetic structure and pH responses of bacterial ASVs across 40 forest soils. (A) Circular phylogenetic tree of all bacterial ASVs detected across the 40 forest sites. Branch colors indicate phylum-level taxonomy (see legend). The outer ring shows the direction of each ASV’s response to soil pH, defined as the sign of the Spearman correlation between ASV relative abundance and soil pH: positive responses indicate increasing relative abundance with higher pH, whereas negative responses indicate decreasing relative abundance with higher pH. The tree is partitioned into 59 response-defined clades identified using PhyloFactor (Kolmogorov–Smirnov tests, p < 0.05), shown as alternating gray and white sectors. (B) Same phylogenetic tree as in (A), with the outer ring indicating the estimated pH optimum of each ASV (range 4.2–6.8). The same clades are shown as alternating gray and white sectors.

We then tested whether response-defined clades improved prediction of pH-driven ASV abundance shifts. We developed a two-stage machine-learning framework that generated predictions at the ASV level. In the first-stage model, ASV-level response values were predicted from response-defined clade identity, ASV identity and TITAN2-derived pH-range features. Including ASV identity allowed the model to account for ASV-specific variation, whereas the response-defined clade term represented shared response structure among related ASVs. This model achieved strong performance (train R² = 0.87; test R² = 0.77; Fig. S6A), with a mean absolute error (MAE) of 14.07 and a mean absolute percentage error (MAPE) of 0.26. Feature-importance analysis indicated that response-defined clade identity contributed strongly to response prediction, followed by ASV identity and pH-range features (Fig. S6B). In the second-stage model, predicted response values were used to forecast ASV-level abundance shifts along pH gradients. This integrated framework achieved high accuracy (train R² = 0.98; test R² = 0.91; test MAE = 2.07; Fig. 4A), with initial abundance and predicted response values being the strongest predictors (Fig. S6C).

**Fig. 4.**
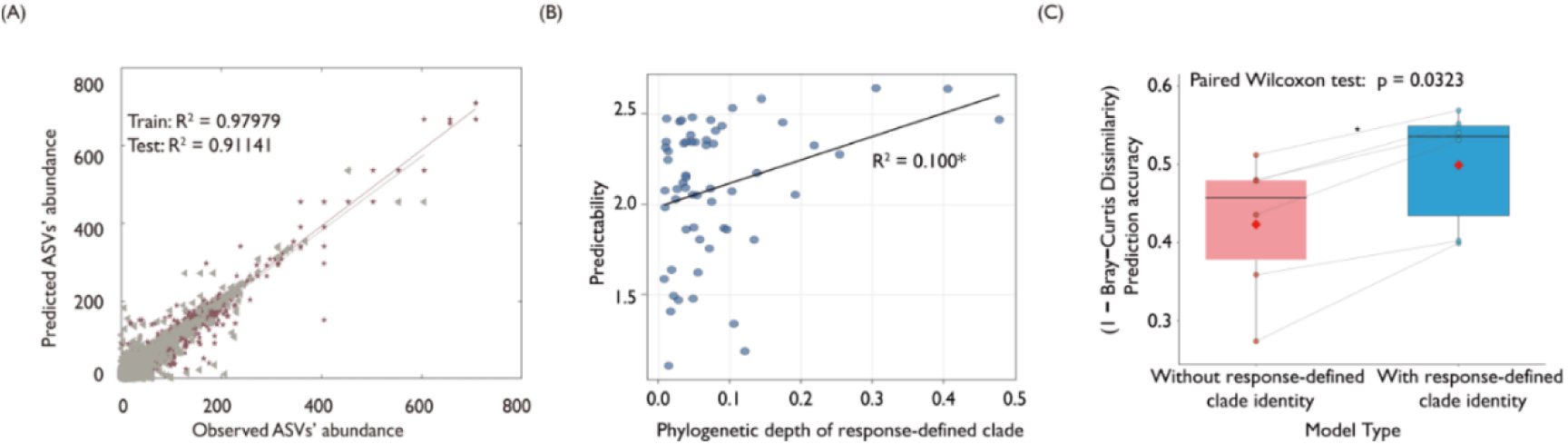
Model performance in predicting ASV abundance shifts under soil pH variation and cross-ecosystem transferability. (A) Relationship between model-predicted and observed ASV abundances across all samples. Purple pentagons represent the training set, and gray triangles represent the test set. The diagonal line indicates the 1:1 relationship (predicted = observed). (B) Relationship between clade-level prediction accuracy and the phylogenetic depth of response-defined clades. Each point represents one clade, and the line shows the fitted linear regression. Statistical significance was assessed using linear regression (p < 0.05). (C) Cross-ecosystem prediction accuracy of bacterial community composition, comparing models with and without response-defined clade identity. For each region, models trained on the remaining 34 forests were used to predict conifer communities from broadleaf communities. Accuracy was defined as 1 − Bray–Curtis dissimilarity between predicted and observed communities. Lines connect paired predictions within each region; statistical significance was assessed using paired Wilcoxon signed-rank tests.

To directly assess the contribution of clade-level response structure to ASV-level predictability, we reran the models without response-defined clade identity while keeping all other features identical. Removing clade identity reduced performance in both stages. For response prediction, test R² decreased from 0.77 to 0.74 and MAE increased from 14.07 to 14.98. For abundance prediction, test R² decreased from 0.91 to 0.87 and MAE increased from 2.07 to 2.38 (Fig. S7). These results indicate that response-defined clades capture shared phylogenetic response information for pH-driven ASV abundance shifts that is not fully explained by ASV identity or range-specific response features alone.

Because our hypothesis further predicts that response-defined clades should be more informative where shared response tendencies persist across greater phylogenetic distances, we next examined whether ASV-level prediction accuracy increased with the phylogenetic depth of response-defined clades. To do so, we summarized ASV-level prediction accuracy within each response-defined clade, yielding a clade-level measure of predictability. Clade-level prediction accuracy increased significantly with phylogenetic depth, although phylogenetic depth explained only a modest fraction of variation in predictability (R² = 0.10, p = 0.015; Fig. 4B). This pattern supports the expectation that response-defined clades provide stronger predictive signals when shared pH-response tendencies persist across greater phylogenetic distances, while also indicating that phylogenetic depth alone does not fully explain predictability. High-predictability clades generally contained more ASVs and showed stronger taxonomic coherence, often being dominated by a single phylum, whereas low-predictability clades tended to include ASVs from multiple phyla (Fig. S8). Together, these analyses based on the 40-site dataset show that response-defined clades capture ecologically relevant structure in pH-driven ASV abundance shifts and that this predictive information tends to be stronger where shared response tendencies persist across greater phylogenetic distances.

### Response-defined clades improve cross-ecosystem prediction of community shifts

As the final step in testing our second hypothesis, we asked whether the predictive information captured by response-defined clades at the ASV and clade levels also scaled up to community-level predictions of pH-driven compositional shifts. We evaluated this using paired broadleaf-conifer forest contrasts across six regions, which provided matched comparisons under similar climate and soil parent material but contrasting soil pH.

In each region, a broadleaf natural forest and an adjacent conifer plantation formed a matched pair under similar climate and soil parent material, with soil pH representing a consistent and major environmental difference between forest types^13^. Broadleaf forests consistently exhibited lower pH than adjacent conifer forests. Using this systematic pH difference, we treated the broadleaf community and its pH as the initial condition and predicted the corresponding conifer community under the observed pH shift. Models were trained using the remaining 34 forests, ensuring that predictions were made for held-out, independent data. Prediction accuracy was quantified as 1 − Bray-Curtis dissimilarity between predicted and observed conifer communities.

Models incorporating response-defined clades achieved significantly higher prediction accuracy than models without clade information (50.0% vs. 42.4%; paired Wilcoxon signed-rank test, p = 0.003; Fig. 4C). Thus, the predictive information captured by response-defined clades improved not only ASV-level abundance predictions in the 40-site dataset, but also community-level predictions in held-out paired forest contrasts. These results support the community-level component of our second hypothesis, indicating that phylogenetically conserved response structure can provide ecologically meaningful information for predicting bacterial community shifts along a pH gradient.

## Discussion

We extended the trait-based phylogenetic framework previously demonstrated within the marine cyanobacterial lineage *Prochlorococcus*^11,12^ to whole soil bacterial communities composed of many coexisting lineages (Fig. 1). We found that environmental factors whose bacterial responses were conserved at deeper phylogenetic levels contributed more strongly to soil bacterial community turnover (Fig. 2), supporting our primary hypothesis. These findings suggest that bacterial biogeographic patterns depend not only on environmental variation itself, but also on the phylogenetic depth at which bacterial responses to that variation are conserved.

The phylogenetic depth of conserved bacterial responses is thought to reflect the physiological complexity and integration of the functional traits underlying those responses^9,10^. In our study, responses to soil pH were conserved at deeper phylogenetic levels than responses to TC, MAT, and MAP (Fig. 2). This may be because pH responses depend on fundamental cellular processes, such as regulation of the proton motive force, membrane lipid remodeling and cytoplasmic pH homeostasis^14–16^. These systems are closely linked to cellular energy metabolism and require coordinated regulation across multiple genes. They may therefore be difficult to reorganize through a small number of mutations or horizontal gene transfer events, leading bacterial pH responses to be conserved at deeper phylogenetic levels. By contrast, responses to other environmental factors, such as soil carbon content, temperature and water availability, may involve more modular and evolutionarily flexible pathways, including membrane desaturases, osmoprotectant systems and carbohydrate-utilization loci^17–19^. Such pathways may be more readily modified over evolutionary time, allowing bacterial responses to these factors to be conserved at shallower phylogenetic levels. The relatively deep conservation of C/N responses may reflect an intermediate case, in which resource-related gradients involve both conserved physiological strategies and more flexible metabolic pathways. This interpretation is consistent with the metagenomic analysis of Liu et al.^13^, which used the same forest dataset and showed that the soil pH gradient was associated with system-wide regulatory integration, including reorganization of sensing-transduction-transcription systems, two-component regulatory systems and coordinated changes in respiratory and nitrogen-regulation pathways. In addition, the differences in phylogenetic depth observed among environmental responses in this study are consistent with experimental manipulation studies showing that soil bacterial responses to pH manipulation by Ca addition are conserved at deeper phylogenetic levels than responses to water-availability, temperature and nutrient-availability manipulations by drought, warming and nitrogen/phosphorus addition, respectively^8^.

Our results further distinguish between two aspects of bacterial pH responses: response direction and pH optimum. Soil pH response direction was conserved at deep phylogenetic levels, whereas pH optima were often not conserved (Fig. 3). This distinction is consistent with previous studies showing that pH optima exhibit only weak phylogenetic signal across bacterial taxa^20^, and that, at biogeographic scales, pH optima are more strongly associated with specific functional genes and metabolic modules than with phylogenetic relatedness^21^. Together, these findings suggest that broad response direction may be constrained by conserved physiological systems, whereas variation in pH optimum may reflect more labile genetic and metabolic features that fine-tune adaptation to local pH conditions. This separation may allow bacterial lineages to maintain conserved pH-response strategies while also adapting to local pH environments.

Phylogenetically conserved bacterial responses were useful not only for explaining community turnover, but also for identifying response-defined clades that function as ecologically meaningful units of community organization, supporting our second hypothesis (Fig. 4). Models using response-defined clades predicted pH-driven ASV abundance shifts more accurately than models that treated ASVs as independent units, indicating that these clades capture shared ecological response information among related ASVs. Moreover, the modest but significant increase in predictability where shared response tendencies persisted across greater phylogenetic distances suggests that phylogenetic depth provides additional, although not exhaustive, information about ASV-level responses. More broadly, our results show that ASV-level responses inferred from biogeographic data can be treated as response trait-like information and mapped onto phylogeny. This allows biogeographic patterns to be interpreted in terms of both taxon-level response traits and the phylogenetic scales at which those responses are conserved, with response-defined clades summarizing shared response tendencies among related ASVs. This view is consistent with the broader perspective that bacterial lineages above the species level can exhibit ecological coherence^22^.

The paired broadleaf-conifer contrasts further showed that response-defined clades improved community-level prediction of pH-driven compositional shifts across ecosystems (Fig. 4). This result extends our second hypothesis from ASV-level responses to community-level shifts, suggesting that pH-linked phylogenetic response structure contains transferable ecological information. Prediction accuracy remained moderate, however, indicating that this response structure captures only part of the community differences between forest types. Differences in litter chemistry, root traits, vegetation history, microbial activities and other biotic or edaphic properties may also contribute to these contrasts^13,23^. Thus, this analysis should be viewed as evidence that response-defined clades improve prediction under realistic paired-forest contrasts, while also highlighting that additional environmental and biotic factors are needed to fully explain forest-type differences.

Together, our findings suggest that the phylogenetic depth of bacterial environmental responses explains why some environmental factors contribute more strongly to community turnover than others. Bacterial biogeographic patterns therefore depend not only on environmental variation itself, but also on the phylogenetic organization of bacterial responses to that variation. Our results further show that phylogenetic clades sharing similar environmental responses can represent ecologically meaningful units for understanding bacterial community responses. Because this study focused on forest ecosystems in Japan, future work should test the generality of this framework across broader biomes and environmental contexts. At the same time, substantial variation remains unexplained, highlighting the need for models that more comprehensively integrate phylogenetic structure, functional traits and context-dependent environmental responses.

## Methods

### Study sites and data source

This study collected data from 40 forest sites across Japan, spanning latitudes from N44°22’ to N26°45’ and longitudes from E128°13’ to E144°39’. Mean annual temperature (MAT) ranged from 4.4 to 20.9 °C, and mean annual precipitation (MAP) ranged from 820 to 3080 mm. The sites encompassed both temperate and subtropical zones and included 15 broadleaved forests and 25 coniferous forests. Mineral soil samples (0–10 cm) were collected from five locations per site during summer or autumn. Soil carbon and nitrogen contents were measured using a CN analyzer, and soil pH was determined by water extraction. Detailed soil properties were published previously^13^.

DNA was extracted from 0.4 g of soil for all 200 samples (five replicates per site across 40 forests). Prokaryotic 16S rRNA genes were amplified using the 515F–806R primers^24^ and sequenced on an Illumina MiSeq platform. The 16S rRNA gene sequencing data are available under DDBJ project PRJDB18602 (accession numbers DRX705571–DRX705769). Sequence processing was performed using UPARSE^25^, VSEARCH v2.22.1^26^, and QIIME2 v2023.2^27^. ASV classification and taxonomic assignment were conducted in QIIME2 using the SILVA database v138^28,29^. The resulting ASV table was rarefied to 22,055 reads per sample using the R package phyloseq^30^. These steps, along with additional analytical details, are documented in Liu et al.^13^. In the present study, we further restricted the dataset to ASVs that could be annotated at the phylum level, were detected in more than 10 samples, and together contributed more than 0.01% of the total reads. This filtering yielded a final dataset of 1,489 ASVs for downstream analyses.

For ASV sequence alignment, we used MAFFT^31^, and phylogenetic tree reconstruction was performed using approximately maximum-likelihood inference implemented in FastTree^32^. Phylogenetic trees were visualized and edited using iTOL^33^.

### Environmental drivers and phylogenetic conservation of bacterial community responses

To test the hypothesis that whether environmental factors whose bacterial responses are conserved at deeper phylogenetic levels contribute more strongly to bacterial community turnover, we combined community-level, phylogenetic, and response-trait-based analyses. First, we used generalized dissimilarity modeling (GDM) to quantify the independent contributions of five environmental variables, soil pH, mean annual temperature (MAT), mean annual precipitation (MAP), soil total carbon (TC), and soil C/N ratio (C/N), to bacterial community turnover across 40 forest sites. Environmental variables were standardized using Euclidean distance, and community dissimilarity was calculated using the Bray-Curtis index. GDM analyses were performed using the gdm package^34^ with backward elimination to retain significant predictors. For each variable, we extracted I-spline curves and deviance contributions, providing directly comparable effect sizes across environmental drivers. Although the workflow followed Liu et al.^13^, models were re-fitted using the selected ASVs and a consistent set of environmental variables for this study.

We next quantified the phylogenetic depth of conserved responses to each environmental factor using consenTRAIT^9^. For each environmental variable, ASV-level responses were first assessed using Spearman correlations between relative abundance and environmental values (p < 0.05; Hmisc package). ASVs were classified as exhibiting positive or negative responses based on the sign of the correlation.

Response-consensus clades were then identified on the phylogenetic tree as clades in which ≥90% of descendant ASVs exhibited a significant response. We defined three response categories: (i) all responses, in which ASVs with significant positive or negative responses were treated as responsive irrespective of direction; (ii) positive responses only; and (iii) negative responses only. The all-responses category tests whether the capacity to respond to an environmental gradient is phylogenetically conserved, irrespective of response direction, such that closely related ASVs may respond in opposite directions yet still share environmental sensitivity. Phylogenetic depth of each response-consensus clade was estimated using the consenTRAIT algorithm with R package castor^35^, and mean depth across clades (τD) was calculated for each environmental variable and response category. To assess phylogenetic conservation, observed τD values were compared with a null distribution generated by permutation of ASV response labels across the phylogeny (1,000 permutations).

We then related the contribution of each environmental variable to community turnover, as estimated by GDM, to the corresponding τD values using linear regression. This analysis tested whether the phylogenetic depth of conserved responses predicted the contribution of each environment factor to bacterial community turnover.

Beyond response direction, we also examined whether response strength was related to phylogeny. Response-strength dissimilarity was calculated as the absolute difference in Spearman correlation coefficients between ASVs. Phylogenetic distances between ASVs were pairwise patristic distances derived from the phylogenetic tree using the R package ape^36^, and the relationship between phylogenetic distance and response-strength dissimilarity was assessed using Mantel tests.

We conducted additional analyses focusing on soil pH. First, we estimated each ASV’s pH optimum, defined as the pH at which its abundance peaks, based on the relative-abundance-weighted mean pH across sampling sites. Second, we examined whether pH responses are phylogenetically conserved within individual bacterial phyla by repeating the response-consensus clade and depth analyses on phylum-specific subtrees. For each phylum, consensus clades were identified and mean phylogenetic depth (τD) was calculated using the same procedure described above.

### Identification of response-defined clades

To test whether phylogenetically conserved responses organize bacterial community shifts and improve their predictability, we focused on soil pH and identified phylogenetically coherent clades that respond consistently to pH, hereafter referred to as response-defined clades. We used PhyloFactor^37^, which applies isometric log-ratio (ILR) transformation to compositional data to identify phylogenetic splits associated with environmental gradients^38^. At each iteration, PhyloFactor tested whether differences in ILR-transformed abundances across phylogenetic partitions were associated with pH using Kolmogoro-Smirnov tests. To ensure robustness, we retained only clades containing ≥5 ASVs, resulting in 59 response-defined clades. For each clade, we calculated its phylogenetic depth, providing a measure of how deeply each pH response is conserved within the phylogeny.

### Quantifying ASV-level pH responses within response-defined clades

Response-defined clades summarize broad phylogenetic response patterns to soil pH, but they may mask nonlinear pH responses and finer-scale variation among ASVs within the same clade. To resolve this within response-defined clade heterogeneity, we quantified ASV-level responses to soil pH within response-defined clades using Threshold Indicator Taxa Analysis (TITAN2^39^). TITAN2 was applied across overlapping sliding windows spanning the full pH range (3.5–8.0), with window widths of 0.2–1.5 pH units (increments of 0.1) and step size of 0.1 pH units.

Within each clade-window, TITAN2 evaluated candidate pH thresholds by partitioning samples at each threshold into low-pH (pH below the threshold) and high-pH (pH at or above the threshold) groups. For each ASV and candidate pH threshold, directional z-scores (sum(z⁺) and sum(z⁻)) and indicator values (IndVal) were calculated, and the threshold maximizing IndVal was selected as the optimal change point. Significance was assessed using 500 bootstrap resamples, retaining only ASVs with purity ≥0.90 and reliability ≥0.90.

From these outputs, we derived three ASV-level metrics: (i) response direction (increasing or decreasing with pH), (ii) response value (absolute IndVal at the optimal threshold), and (iii) effective pH range, defined as the set of windows in which the ASV exhibited a consistent and significant response. This procedure provides a clade-aware characterization of ASV responses across the pH gradient.

### Two-stage ensemble learning for predicting bacterial abundance under pH shifts

To test whether response-defined clades improve pH-driven ASV abundance shifts, we developed a two-stage ensemble learning framework that integrates response-defined clades with ASV-level response metrics. The framework consists of two linked models: Model I predicts pH response values, and Model II predicts final bacterial abundance under altered pH conditions. The workflow follows a supervised-learning structure with paired input-output data (Fig. S9).

Model I predicts ASV-level pH responses using five input features: (1) response-defined clade identity (categorical identifiers for the 59 response-defined clades representing a group of ASVs sharing similar responses), (2) ASV identity capturing the ASV-specific response, (3) initial soil pH (pH_left_), (4) target soil pH (pH_Right_), and (5) pH_Range_ (difference between upper and lower effective pH limits) (Fig.S9). predicts ASV abundance under the target pH using seven input features: (1) response-defined clade identity, (2) ASV identity, (3) pH_left_, (4) pH_Right_, (5) pH_Range_, (6) initial abundance at pH_left_, and (7) predicted response values from Model I (Fig.S9).

To leverage complementary model structures, we implemented a hybrid ensemble combining a multilayer perceptron (MLP) and a decision tree (DT). The MLP consisted of two hidden layers (16 neurons each) with ReLU activation functions to capture nonlinear relationships between input features and pH response values. Dropout regularization (rate = 0.3) was applied to reduce overfitting, and the Adam optimizer was used for parameter estimation. The DT model used mean squared error (MSE) as the splitting criterion and was used to model the regression problem of predicting abundance changes. Both models were trained and fine-tuned on the same dataset with the goal of minimizing prediction error. Final predictions were obtained using a threshold-based ensemble rule: when the difference in validation error between models was below θ = 0.05, predictions were combined with weights inversely proportional to validation error; otherwise, the better-performing model was selected (Fig. S9).

To ensure robust model evaluation, the dataset was partitioned into training (80%), validation (10%), and test (10%) sets based on response-defined clades. Each subset contained a comparable representation of all response-defined clades, preserving clade representation across splits.

To directly assess the contribution of clade-level response structure to ASV-level predictability, we compared model performance with and without inclusion of response-defined clade identity. In the full model, clade identity was included as a categorical predictor representing groups of ASVs with similar pH responses, allowing the model to capture shared response patterns among related taxa. In the reduced model, clade identity was excluded, and predictions were based on ASV identity and environmental variables alone. Specifically, Model I predicted ASV response values using ASV identity, pH_left_, pH_Right_, and pH_Range_, and Model II predicted ASV abundance under target pH conditions using predicted response values together with ASV identity, pH_left_, pH_Right_, and pH_Range_, and initial abundance. Both models were independently optimized prior to comparison. Improvement in predictive performance when incorporating clade identity indicates that phylogenetically conserved responses contribute to predictable community reorganization under soil pH change.

To further interpret the relative importance of clade-level response structure within the modeling framework, we quantified feature importance for both Model I and Model II. Feature importance was assessed based on the relative contribution of each predictor to model performance, allowing evaluation of the role of response-defined clade identity in shaping predictions.

### Prediction accuracy assessment of response-defined clades

To evaluate whether predictive accuracy varies with the phylogenetic depth of conserved response, we assessed clade-level prediction performance using three complementary metrics: coefficient of determination (R²), mean absolute error (MAE), and mean absolute percentage error (MAPE). These metrics capture complementary aspects of model performance. R² evaluates how well the model explains the variance in the data, MAE measures the absolute magnitude of the prediction error, and MAPE evaluates the relative error. These metrics were calculated for each clade based on predictions across its constituent ASVs.

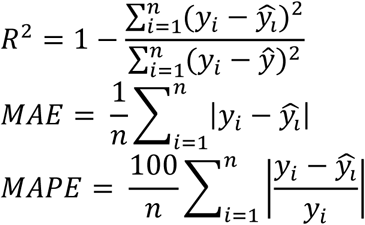

where yi is the actual value, ŷi is the predicted value, ŷ is the mean of actual values, and n is the number of ASVs in each clade.

To obtain a unified measure of prediction accuracy, all metrics were scaled to a range of 0–1 using min–max normalization. MAE and MAPE were inverted so that higher values indicate better performance. The composite accuracy score for each clade was then calculated as the mean of the normalized metrics:

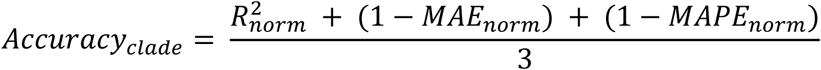

We then assessed the relationship between clade-level prediction accuracy and phylogenetic depth using linear regression.

To identify taxonomic patterns associated with predictability, we selected the top and bottom 10 clades based on accuracy scores. These clades were then mapped to their corresponding phylum- and genus-level classifications.

### Predicting bacterial community shifts across forest types

To test whether the predictive information captured by response-defined clades at the ASV and clade levels also scaled up to community-level predictions of pH-driven compositional shifts, we performed a cross-ecosystem transfer test across six regions, each consisting of a matched pair of a broadleaf natural forest and an adjacent conifer plantation. Climate, soil type, and soil C and N contents were similar between forest types, whereas soil pH consistently differed, with broadleaf forests exhibiting lower pH than conifer forests^40^. These paired sites provided a natural setting in which vegetation-driven pH change was the dominant environmental shift.

To ensure rigorous independent validation, we applied the following procedure. For each of the six target forest sites, all conifer forest samples (five replicates per site; 30 samples in total) were excluded from the dataset. TITAN2 analyses were then re-run using the remaining samples (n = 170). Both Model I and Model II were subsequently retrained, with hyperparameters independently re-optimized for each of the two frameworks (“with clade identity” and “without clade identity”), as described above. The retrained models were then used to predict how the broadleaf forest community at each target site would shift under the soil pH conditions of the corresponding conifer forest.

Specifically, Model I used the initial soil pH of the broadleaf forest (pH_left_) and the target soil pH of the conifer forest (pH_right_), together with their difference (pH_range_) and ASV identity, as input features. In the “with clade identity” framework, clade identity was included as an additional feature, whereas it was excluded in the “without clade identity” framework. The output of this stage was a predicted pH response value for each ASV. In the second stage, Model II used these predicted responses to estimate ASV abundances under the soil pH conditions of the conifer forest (i.e., target abundances at pH_right_). Input features included pH_left_, pH_right_, pH_range_, ASV identity, and the initial abundance at pH_left_ derived from the broadleaf forest community. Initial abundance was defined at the regional level by averaging ASV relative abundances across the five broadleaf replicates, yielding one community per region. Observed conifer communities (target communities) were defined analogously. Soil pH values (pH_left_ and pH_right_) were also averaged across replicates within each forest type. As in the first stage, clade identity was included only in the “with clade identity” framework.

Predicted communities were compared with the observed conifer communities, which were defined as regional means by averaging ASV relative abundances across the five conifer replicates, using Bray-Curtis dissimilarity. Prediction accuracy was quantified for each region as 1 – Bray-Curtis dissimilarity between predicted and observed conifer communities, and differences the two frameworks were evaluated using paired Wilcoxon signed-rank tests.

## Acknowledgements

This work was financially supported by National Key R&D Programmes of China (Grant Number 2022YFF0801801), Beijing Municipal Natural Science Foundation (Grant Number IS23072), Research Fund for International Scientists of National Natural Science Foundation of China (Grant Number 32350610251).

---

**Table S1.** Mean phylogenetic depth (τD) of response-consensus clades and permutation-based tests of phylogenetic conservation across environmental variable.

| Environmental variable | All response | Positive response | Negative response |
| --- | --- | --- | --- |
| pH | **0.023** | **0.019** | **0.015** |
| C/N | **0.017** | **0.013** | **0.019** |
| MAT | **0.013** | 0.008 | **0.010** |
| TC | 0.012 | 0.01 | **0.011** |
| MAP | 0.011 | 0.008 | **0.009** |

Response categories include all significant responses irrespective of direction (“All”), positive responses only (“Positive”), and negative responses only (“Negative”). Phylogenetic depth of each clade was estimated using the consenTRAIT algorithm, and τD represents the mean phylogenetic depth across response-consensus clades. Phylogenetic conservation was assessed by comparing observed τD values with a null distribution generated by permutation of ASV responses across the phylogeny; Bold values indicate significant deviation from the null expectation ( $p < 0.05$ ).

**Table S2.** Mean phylogenetic depth (τD) of response-consensus clades based on soil pH responses and permutation-based tests of phylogenetic conservation across major bacterial phyla.

| Phylum | Number of ASVs | All response | Positive response | Negative response |
| --- | --- | --- | --- | --- |
| Planctomycetota | 44 | **0.043** | **0.031** | **0.058** |
| Chloroflexi | 49 | 0.035 | 0.023 | 0.049 |
| Gemmatimonadota | 20 | 0.028 | 0.032 | 0.034 |
| Myxococcota | 44 | **0.023** | **0.032** | **0.014** |
| Bacteroidota | 26 | 0.023 | **0.031** | 0.005 |
| RCP2-54 | 31 | 0.016 | 0.061 | 0.011 |
| Actinobacteriota | 115 | **0.015** | **0.016** | **0.019** |
| Verrucomicrobiota | 98 | **0.015** | **0.015** | **0.017** |
| Proteobacteria | 458 | **0.011** | **0.014** | **0.010** |
| Acidobacteriota | 493 | **0.006** | **0.011** | **0.005** |

**Fig S1.**
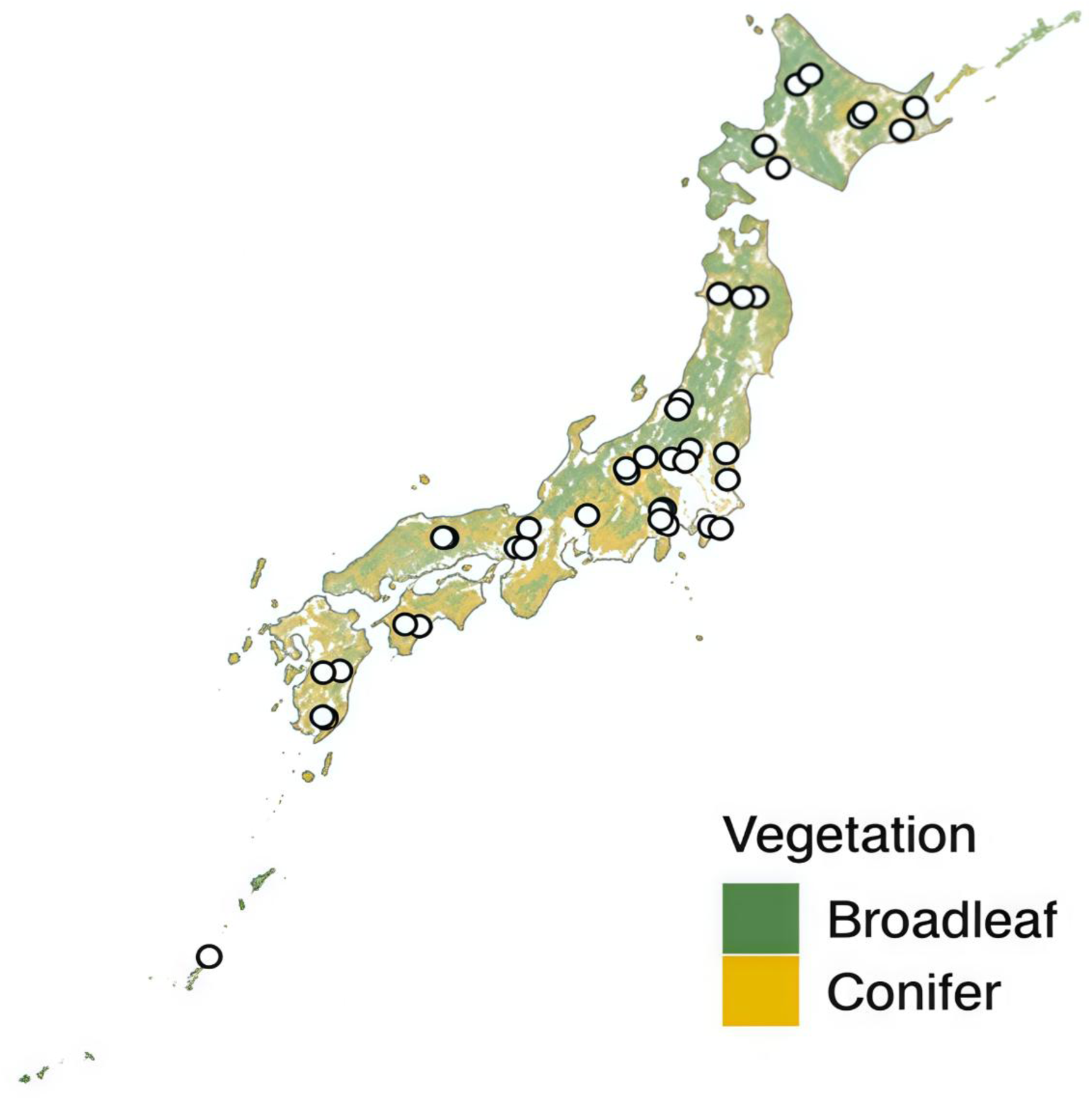
Geographic distribution of the 40 forest soil sampling sites across Japan. Soil bacterial communities were sampled from 40 forest sites spanning broad environmental gradients, including soil pH, mean annual temperature (MAT), mean annual precipitation (MAP), soil total carbon (TC), and soil C/N ratio. Background colors indicate dominant forest vegetation type, with green representing broadleaf forests and yellow representing coniferous forests.

**Fig S2.**
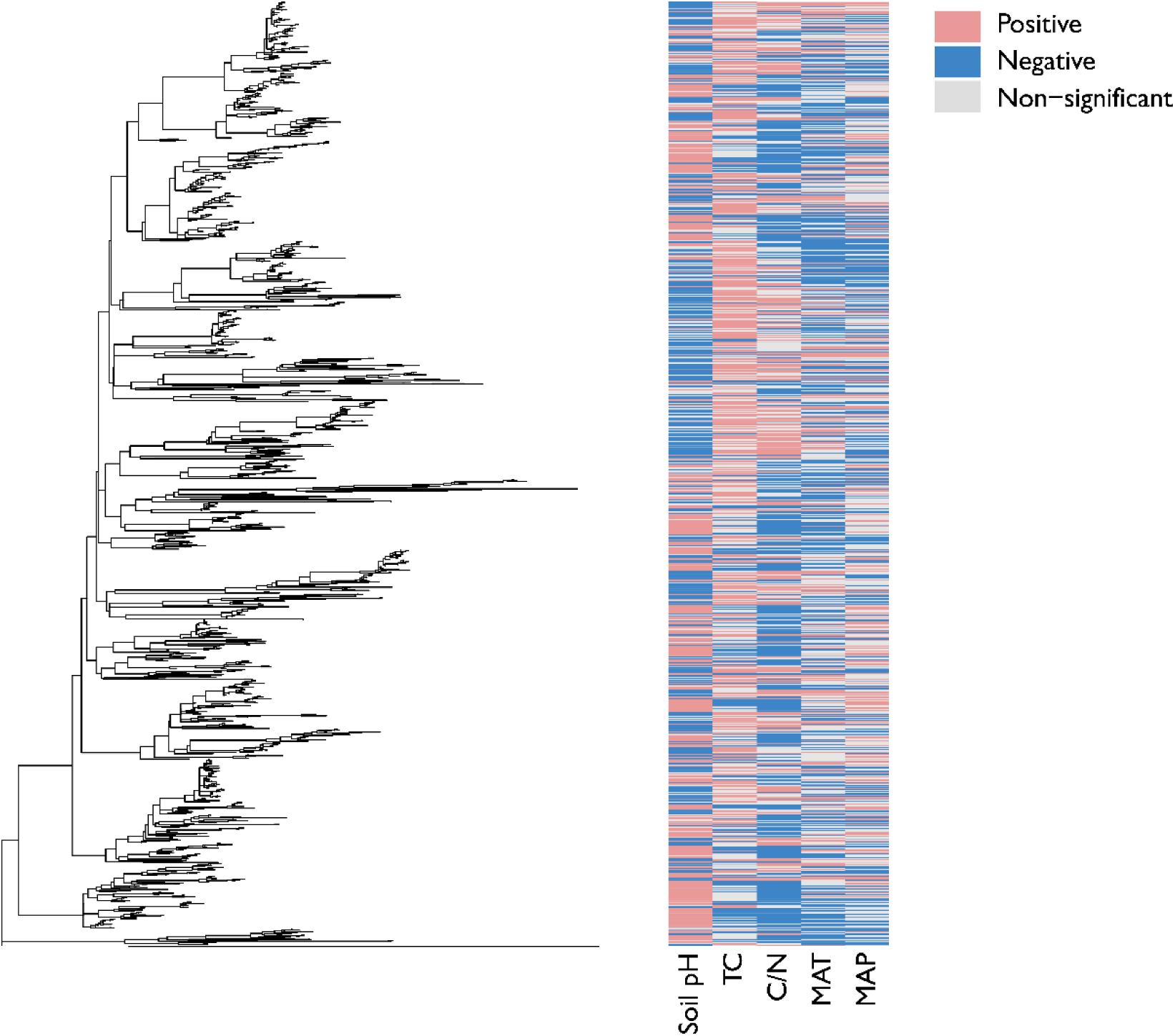
Phylogenetic distribution and direction of ASV responses to multiple environmental factors. The phylogenetic tree (left) illustrates evolutionary relationships among all detected ASVs. The adjacent heatmap (right) summarizes the direction of each ASV’s abundance response to five major environmental gradients: soil pH, total carbon (TC), C:N ratio (C/N), mean annual temperature (MAT), and mean annual precipitation (MAP). For each ASV–environment pair, Spearman correlation coefficients (ρ) were calculated between ASV relative abundance and the corresponding environmental variable across all sites. Cells are color-coded by the sign of ρ and P-value: pink indicates a significant positive correlation (ρ > 0, P < 0.05), implying higher ASV abundance with increasing environmental values (e.g., higher pH or MAT); blue indicates a negative correlation (ρ < 0, P < 0.05), implying decreasing abundance along that gradient; Gray indicate non-significant correlations.

**Fig S3.**
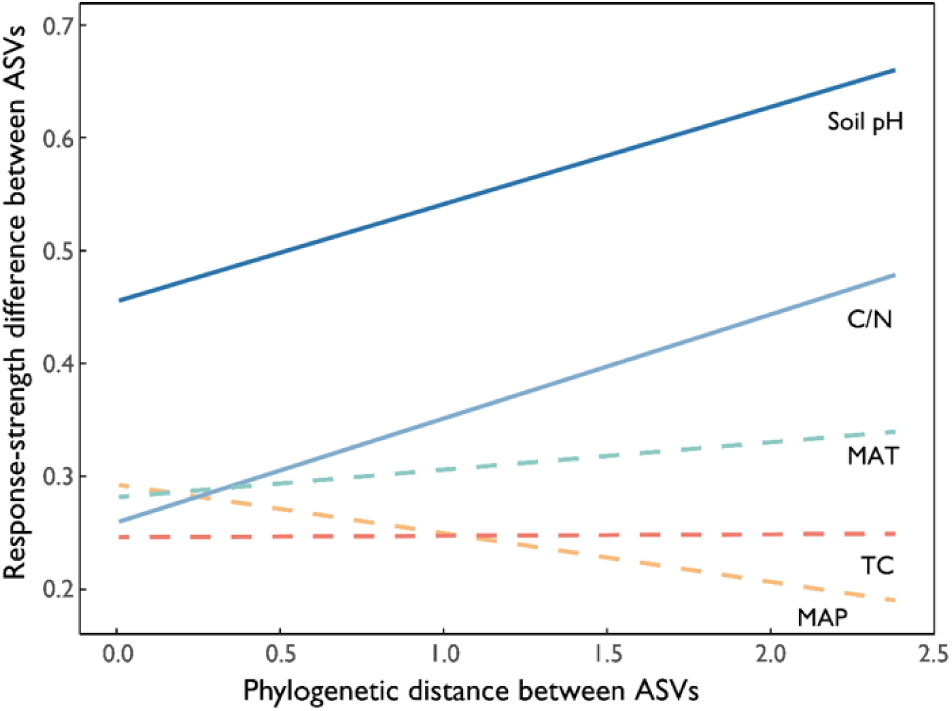
Mantel test results showing relationships between phylogenetic distance and response-strength difference of bacterial ASVs to environmental factors. Response-strength difference was calculated as the absolute difference in Spearman correlation coefficients (ρ) between two ASVs, where ρ is the correlation between ASV relative abundance and each environmental variable. Phylogenetic distances between two ASVs was measured based on branch length between them. Solid lines indicate significant Mantel correlations (P < 0.05), whereas dashed lines indicate non-significant relationships (P ≥ 0.05).

**Fig. S4.**
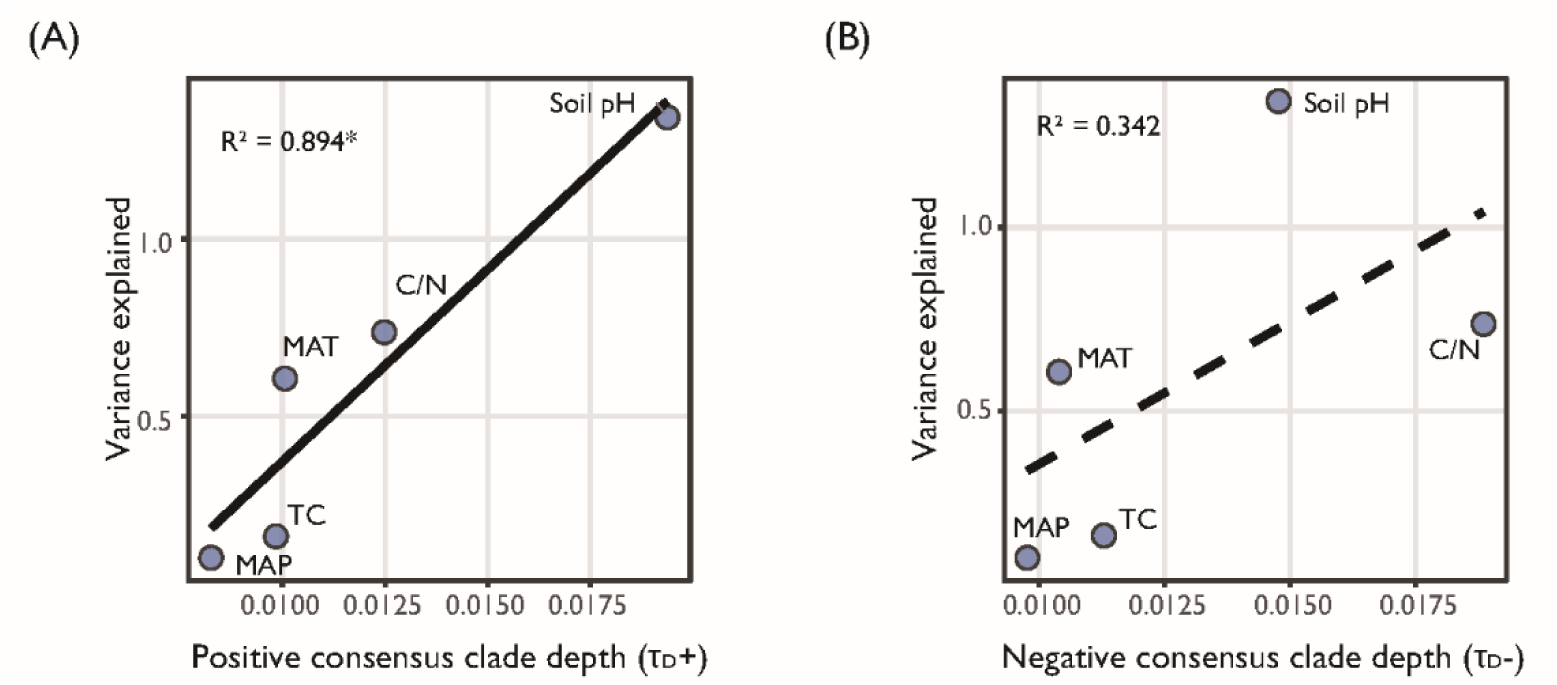
Relationship between the mean phylogenetic depth (τD) of response-consensus clades and the contribution of environmental variables to community turnover, separated by response direction. Response-consensus clades were defined as clades in which ≥90% of descendant ASVs exhibit a significant response (Spearman correlation, p < 0.05), and phylogenetic depth was estimated using the consenTRAIT algorithm. τD represents the mean phylogenetic depth across response-consensus clades and was calculated separately for clades showing positive (A) and negative (B) responses to each environmental variable. The x-axis denotes τD for positive or negative responses, and the y-axis denotes the contribution of each environmental variable to community turnover as derived from the generalized dissimilarity model (GDM; Fig. 2A). A significant relationship was observed for positive responses (A; p < 0.01), whereas no significant relationship was detected for negative responses (B; p ≥ 0.29).

**Fig. S5.**
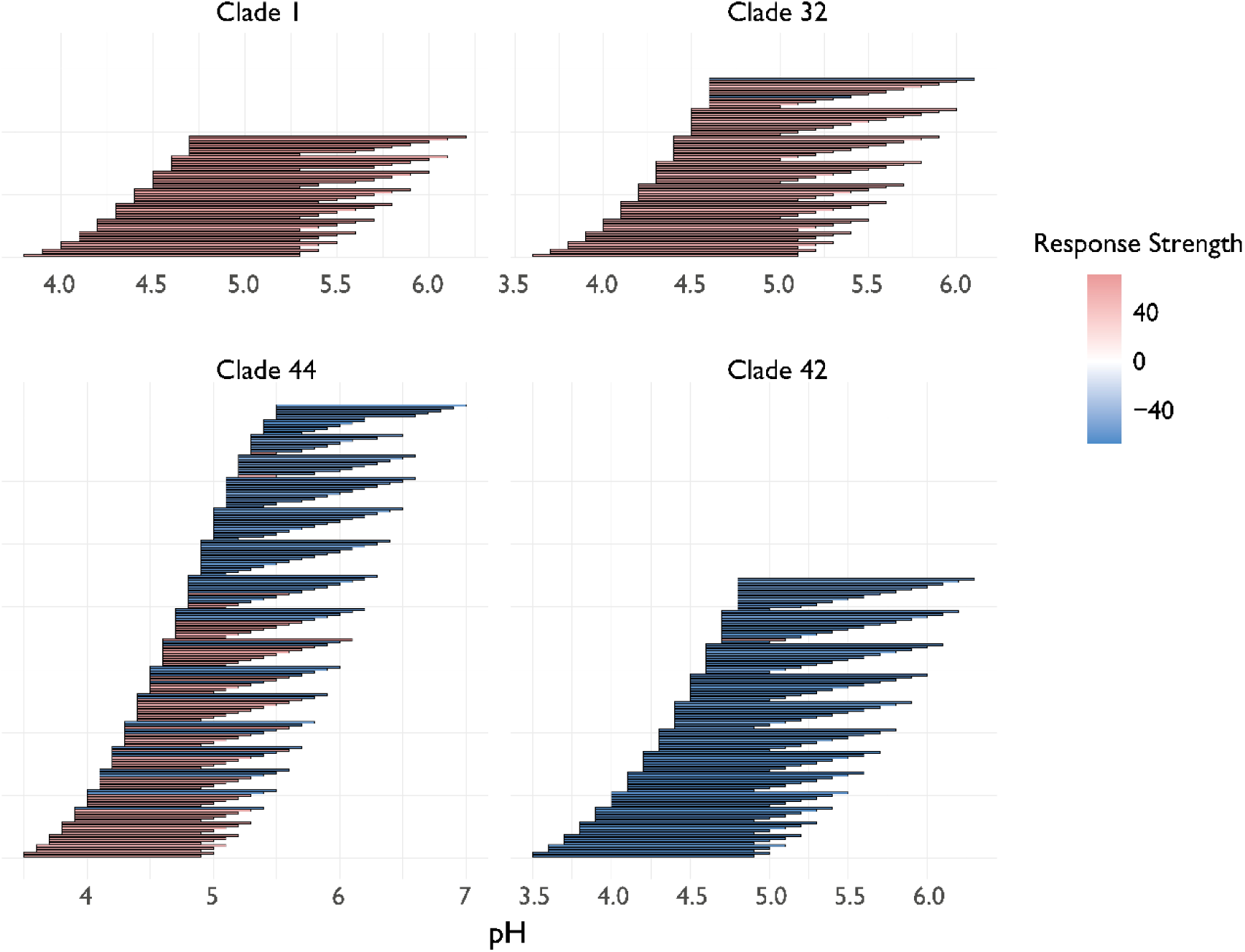
Representative pH response profiles across selected response-defined clades based on TITAN2 analysis. Each horizontal bar represents a sliding window along the pH gradient, with endpoints indicating the lower and upper pH boundaries of the window. Bar color indicates the dominant response direction of ASVs within each clade for that window, based on TITAN2 z-scores: red indicates a positive response (increasing abundance with higher pH; z⁺ > 0), and blue indicates a negative response (decreasing abundance with higher pH; z⁻ < 0). Response strength reflects the mean standardized indicator value (IndVal) of ASVs within each clade. The four clades shown (Clades 1, 32, 42, and 44) illustrate contrasting patterns of pH responses: Clades 1 and 32 are dominated by positive (high-pH) responses, Clade 42 by negative (low-pH) responses, and Clade 44 exhibits mixed responses across the pH gradient.

**Fig. S6.**
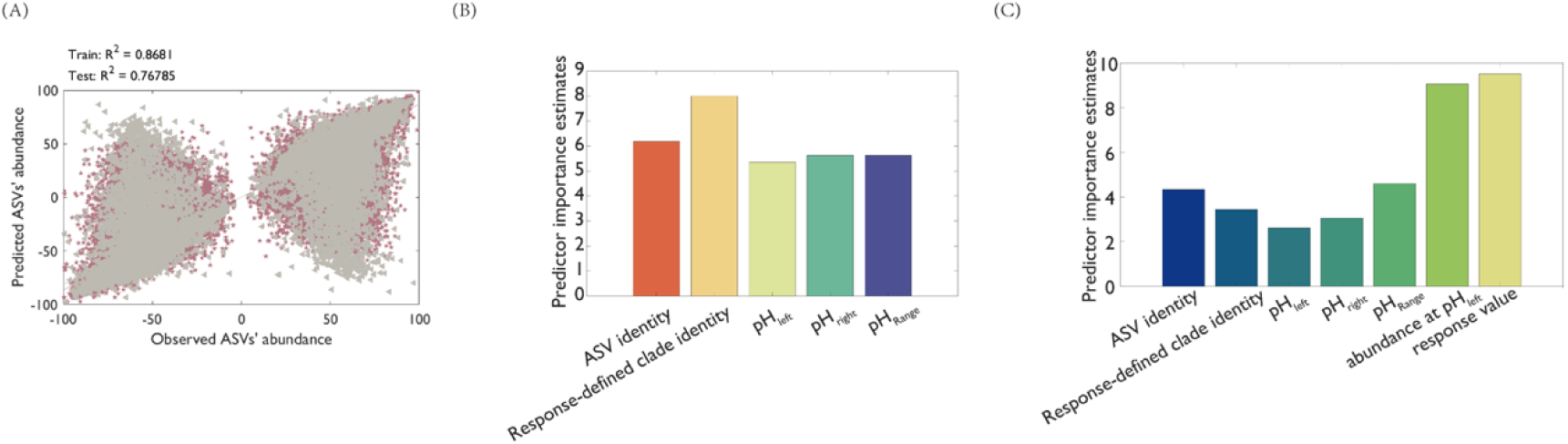
Performance and feature importance of the two-stage ensemble model. (A) Relationship between observed and model-predicted pH response values for ASVs across response-defined clades and sliding pH windows. Observed values (x-axis) are TITAN2-derived response values (standardized IndVal; −99.7 to 99.0), and predicted values (y-axis) are outputs from the Stage I model. Each point represents an ASV–sample combination; purple points indicate the training set and grey points indicate the independent test set. The diagonal line denotes the 1:1 relationship. The model achieved R² = 0.868 for the training data and R² = 0.768 for the test data. (B) Feature importance for the Stage I model predicting ASV-level pH response values. (C) Feature importance for the Stage II model predicting ASV abundance under shifted soil pH conditions.

**Fig. S7.**
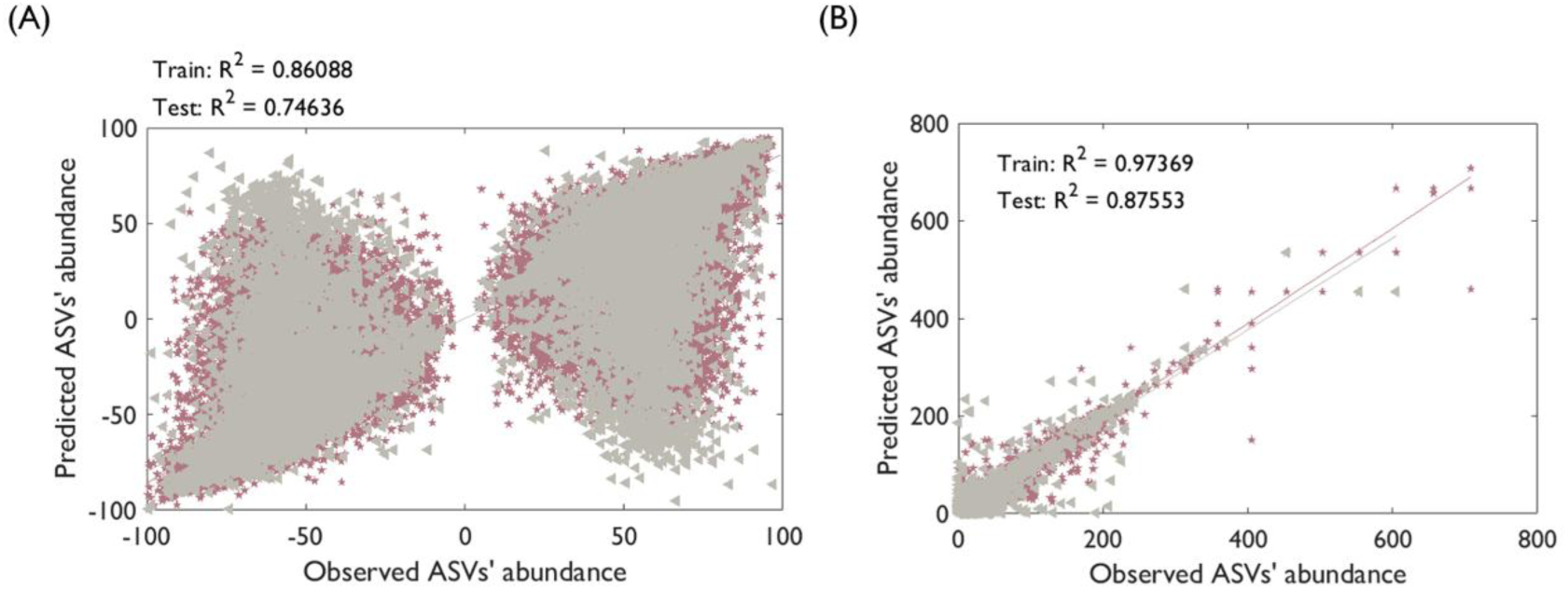
Performance of the two-stage ensemble model excluding response-defined clade identity. (A) Relationship between observed and predicted ASV-level pH response values when response-defined clade identity was excluded from the Stage I model. Observed values (x-axis) are TITAN2-derived response values (standardized IndVal; −99.7 to 99.0), and predicted values (y-axis) are model outputs. Each point represents an ASV–sample combination; purple points indicate the training set and grey points indicate the independent test set. The diagonal line denotes the 1:1 relationship. Compared with the full model (Fig. S6A), removal of clade identity slightly reduced predictive performance (R² = 0.861 for training and R² = 0.746 for testing), indicating that incorporating response-defined clade identity improves model accuracy. (B) Relationship between observed and predicted ASV abundances under target pH conditions for the Stage II model excluding response-defined clade identity. Observed values (x-axis) are measured ASV abundances, and predicted values (y-axis) are model outputs. Each point represents an ASV–sample combination; purple points indicate the training set and grey points indicate the test set. The diagonal line denotes the 1:1 relationship. The model achieved R² = 0.974 for the training data and R² = 0.876 for the test data; the corresponding model including response-defined clade identity is shown in Fig. 4A.

**Fig S8.**
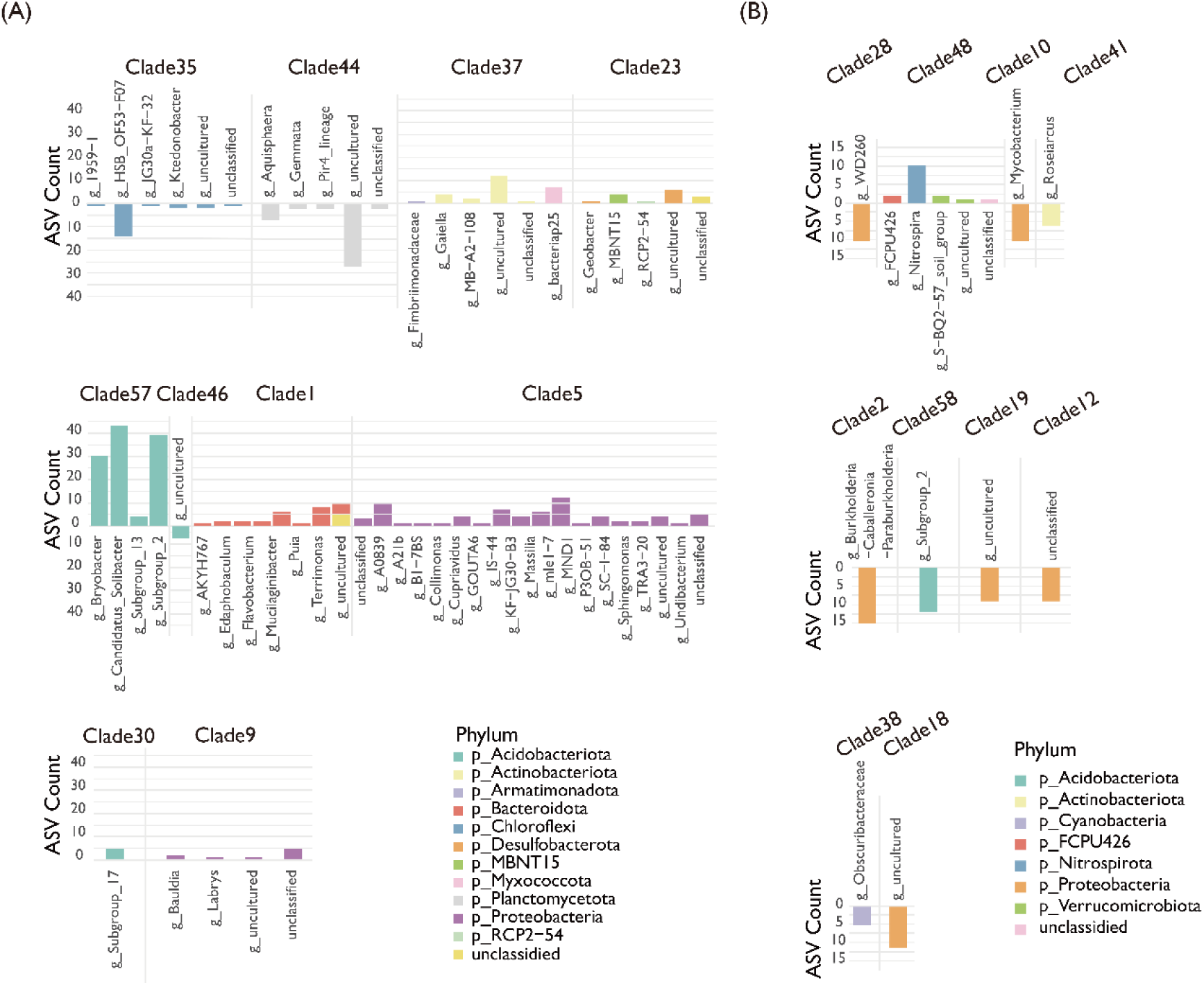
Taxonomic structure of response-defined clades with contrasting predictability. (A) Taxonomic composition of the 10 clades with the highest abundance-prediction accuracy (high-predictability clades). (B) Taxonomic composition of the 10 clades with the lowest abundance-prediction accuracy (low-predictability clades). In both panels, stacked bars show genus- and phylum-level composition for each clade. Colors indicate phylum assignment. The upper part of each bar represents the number of ASVs with positive pH responses, and the lower part represents the number of ASVs with negative pH responses.

**Fig S9.**
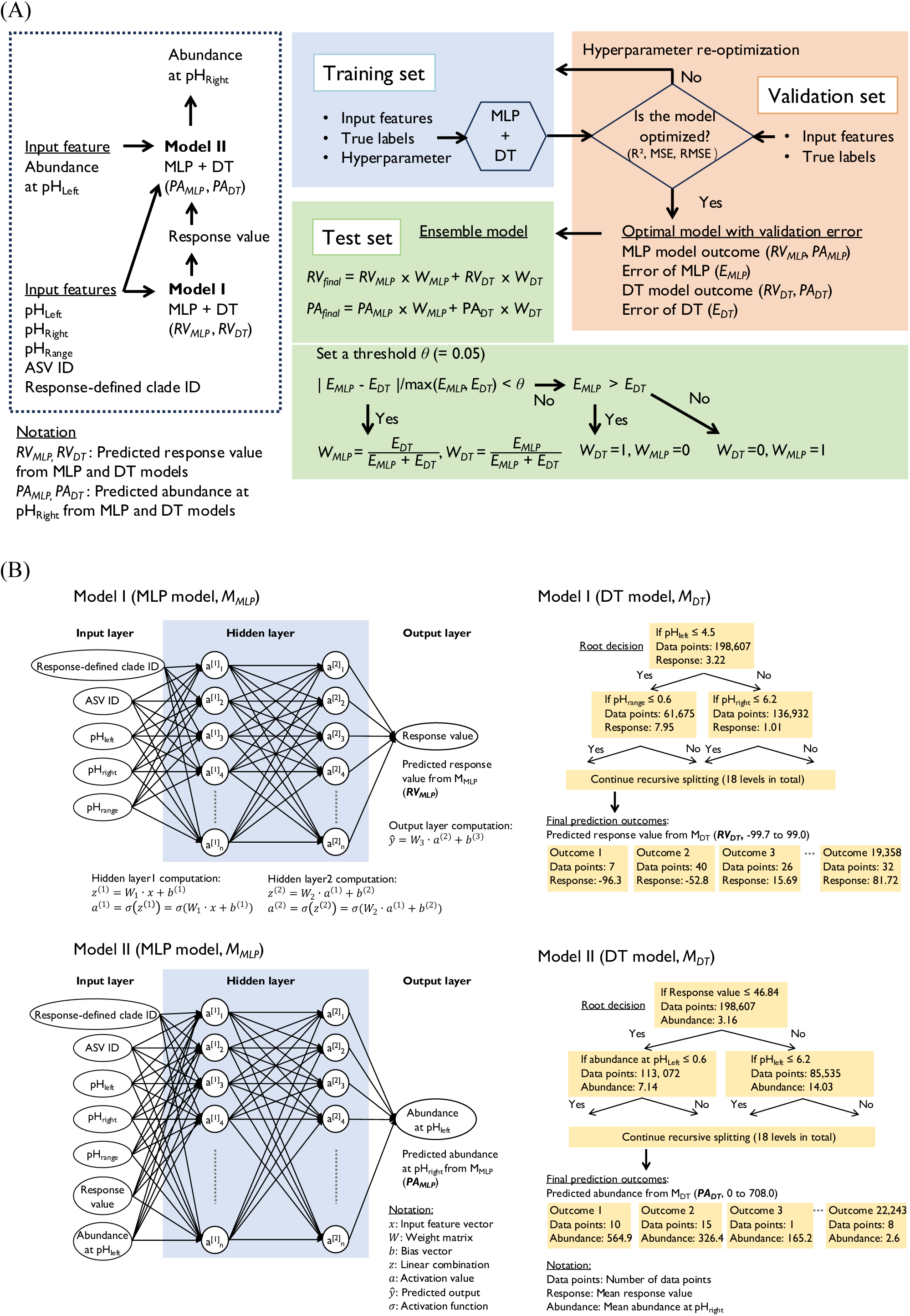
Two-stage ensemble learning framework for predicting microbial abundance responses to pH variation. **(A) Overall architecture of the two-stage ensemble framework.** The framework predicts ASVs’ abundance at target soil pH (pH_right_) through a two-stage process: Stage I estimates the response value (magnitude and direction of abundance change) from response-defined clade identity, ASV identity, and pH descriptors (pH_left_, pH_right_, pH_range_); Stage II then integrates this predicted response with ASV’s abundance at initial soil pH (pH_left_) to forecast the final abundance. Training proceeds on a Training Set consistin of Input Features, True Labels, and a Hyperparameter Search Space. Two algorithms, Multi-Layer Perceptron (MLP) and Decision Tree (DT), are trained in parallel, and their Validation Errors (E_MLP_, E_DT_) are evaluated on a Validation Set. If the normalized difference | E_MLP_ *–* E_DT_ | / max (E_MLP_, E_DT_) is below the threshold θ = 0.05, an Ensemble Model combines both predictions with inverse-error weights (W_MLP_, W_DT_). Otherwise, the model with the smaller validation error is selected as the Optimal Model. Final predictions are evaluated on a Test Set containing unseen samples. **(B) Structure of stage I (Model I) and stage II (Model II).** The framework consists of Stage I (top panels) and Stage II (bottom panels), each combining a Multi-Layer Perceptron (MLP) and Decision Tree (DT) in parallel. Stage I (Model I, M_MLP_ and M_DT_) predicts pH response values from five inputs: response-defined clade identity, ASV identity, pH_left_, pH_right_, and pH_range_. The MLP component (top left) contains an input layer, two hidden layers with 16 neurons each (ReLU activation, dropout = 0.3), and outputs the response value through weighted connections (W₁, W₂, W₃) and biases (b¹, b², b³) optimized via Adam algorithm. The DT component (top right) recursively partitions the feature space across 18 levels based on thresholds (e.g., pH_left_ ≤ 4.5, pH_range_ ≤ 0.6), generating thousands of leaf nodes with predicted response values ranging from −99.7 to 99.0. Stage II (Model II, M_MLP_ and M_DT_) predicts ASV abundance at the target pH using seven inputs: the five features from Stage I plus the abundance at pH_left_ and the predicted response value from Stage I. The MLP architecture (bottom left) mirrors Stage I but outputs abundance at pH_right_. The DT (bottom right) similarly partitions data into leaf nodes with predicted abundances ranging from 0 to 708.0, using splits such as Response value ≤ 46.84 and Abundance at pH_left_ ≤ 0.6. Together, these two stages transform taxonomic and environmental features into accurate quantitative predictions of how each ASV’s abundance changes under new pH conditions.

